# Chronic semaglutide treatment enhances the incentive motivational value of a small food reward and associated cue in male and female rats

**DOI:** 10.64898/2025.12.06.692775

**Authors:** Stephen E. Chang, Christopher A. Turner, Natalia Morales Pagán, Daniela Pereira, Sophia Kleer, Shelly B. Flagel

## Abstract

**Rationale:** Glucagon-like peptide-1 (GLP-1) receptor agonists, such as semaglutide, are increasingly utilized in clinical practice due to their efficacy in promoting sustained weight loss following chronic administration. While acute treatment with GLP-1 receptor agonists has been shown to suppress food intake and reward-seeking behaviors in rodent models, the impact of prolonged exposure on preclinical measures of motivated behavior remains insufficiently characterized.

**Objectives:** This study aimed to systematically evaluate the effects of chronic administration of semaglutide on both the acquisition and expression phases of Pavlovian conditioned approach (PavCA)—a behavioral paradigm used to assess the attribution of incentive salience to a food-paired cue. The influence of chronic semaglutide on the conditioned reinforcing properties of the food-associated cue, performance on a progressive ratio (PR) schedule for food reward, and ad libitum consumption of the food reward were also assessed.

**Results:** Chronic semaglutide administration did not significantly alter either the acquisition or the expression of PavCA behavior. However, relative to vehicle-treated controls, semaglutide markedly enhanced responding for the food-associated cue during a conditioned reinforcement test and increased PR responding for the food reward. In contrast, semaglutide reduced both free consumption of the food reward and homecage chow intake.

**Conclusions:** These findings demonstrate that chronic semaglutide administration potentiates the incentive value of food-paired cues and increases motivation for food reward under restricted access conditions, yet attenuates overall food consumption when food is freely available. This dissociation highlights the nuanced effects of semaglutide on motivated behavior and suggests an amplification of the reinforcing properties of discrete, limited food rewards and associated cues.

## Introduction

Although initially designed to treat type-2 diabetes, glucagon-like-peptide-1 (GLP-1) receptor agonists, such as semaglutide, have surged in popularity due to their ability to induce and maintain weight loss (Wilding et al., 2021; Davies et al., 2021; Wadden et al., 2021; Rubino et al., 2021; Garvey et al., 2022). In humans, semaglutide has been demonstrated to induce an average 11.85% reduction in baseline weight among obese individuals without type-2 diabetes (Tan et al., 2022; Qin et al., 2024). This weight loss has been attributed to semaglutide’s ability to lower food intake, diminish hunger, decrease preference for high-fat foods, and reduce cravings (Blundell et al., 2017). Given the benefits, subcutaneous once-weekly semaglutide administration was approved for weight loss by the US Food and Drug Administration (FDA) in 2021 (Chao et al., 2023).

Studies with rodents have shown that GLP-1 receptor agonists can modify motivated behavior for both food and drug rewards. In the context of food motivation, GLP-1 receptor agonists reduce food intake, conditioned place preference for previously preferred chocolate pellets, and progressive ratio responding for sucrose pellets (Raun et al., 2007; Dickson et al., 2012; Richard et al., 2016; Colvin et al., 2020; Ghidewon et al., 2022; Wharton et al., 2023). Similarly, GLP-1 receptor agonists reduce consumption of alcohol (Vallöf et al., 2016; Colvin et al., 2020; Aranäs et al., 2023), attenuate self-administration of opioids (Zhang et al., 2021; Douton et al., 2022a,b) and cocaine (Schmidt et al., 2016; Hernandez et al., 2018), and diminish relapse-like behavior for each of these drug classes. Thus, there is ample evidence from preclinical studies demonstrating the involvement of GLP-1 signaling in food motivation and addiction-related behaviors.

A psychological process that drives both food- and drug-seeking behaviors is incentive salience attribution (Berridge, 1996; Berridge, 2001). When environmental cues previously paired with rewards are attributed with incentive value, they are transformed into “motivational magnets” (Berridge, 2004), acquiring the ability to capture attention and control reward-seeking behavior, sometimes to an inordinate degree (Berridge & Robinson, 2003). We postulated that GLP-1 receptor agonists may exert their effects on motivated behavior by modulating incentive salience attribution. To test this, we employed a Pavlovian conditioned approach (PavCA) paradigm, in which the insertion of a lever-cue (conditioned stimulus, CS) is paired with the delivery of a food reward (unconditioned stimulus, US). Once the cue-reward association is learned, some rats will approach, press, and bite the lever-cue, reflecting the attribution of incentive salience. This conditioned response is known as sign-tracking (Brown & Jenkins, 1968; Hearst & Jenkins, 1974). Other rats, upon lever-cue presentation, will direct their behavior towards the location of impending reward delivery (i.e., the food cup) rather than the cue itself (Boakes et al., 1978). This conditioned response is known as goal-tracking. Both sign-trackers (i.e., rats that primarily sign-track) and goal-trackers (i.e., rats that primarily goal-track) place predictive value on the lever-cue, but sign-trackers also attribute incentive value to the cue (Flagel & Robinson, 2017). This is evident not only in their PavCA behavior, but also in their performance on a conditioned reinforcement test. Specifically, sign-trackers show increased responding for the lever-cue even in the absence of the food reward (Flagel & Robinson, 2017).

The present set of experiments investigated the effects of chronic semaglutide administration on the acquisition and expression of PavCA behavior—that is, the tendency to sign-track or goal-track—as well as the conditioned reinforcing properties of a food-cue. Progressive ratio responding and free consumption of a food reward were also investigated. Given prior work showing that GLP-1 receptor agonists reduce motivation for food and drug rewards (Trapp & Brierley, 2021; Klausen et al., 2022), we hypothesized that, relative to vehicle, chronic semaglutide administration would reduce sign-tracking behavior, attenuate the conditioned reinforcing properties of the food cue, and diminish progressive ratio responding and free consumption of the food reward.

## Methods

All procedures conformed to *The Guide for the Care and Use of Laboratory Animals* (8^th^ edition, National Research Council, 2011) and were approved by the University of Michigan Institutional Animal Care and Use Committee.

The experimental timelines for each of three experiments that were conducted to assess the effects of chronic semaglutide on PavCA behavior, the conditioned reinforcing properties of the reward cue, progressive ratio responding for the food reward, and homecage and free-feeding consumption are shown in **Figure 1**. In Experiment 1, semaglutide was given prior to and during the acquisition of PavCA behavior. In Experiment 2, semaglutide was given after rats had acquired PavCA behavior and during tests for the expression of PavCA behavior. In both Experiments 1 and 2, following PavCA sessions, the effects of semaglutide on the conditioned reinforcing properties of the food-associated cue and progressive ratio responding for the food reward were assessed. In Experiment 3, the effects of chronic semaglutide were investigated on homecage chow consumption and free consumption of the food reward used in Experiments 1 and 2. For all experiments, the effects of chronic semaglutide on locomotor response to novelty (i.e., “sensation seeking”) was assessed following the last behavioral assay (see Supplementary Information for Methods and Results pertaining to locomotor response to novelty).

**Figure 1.**
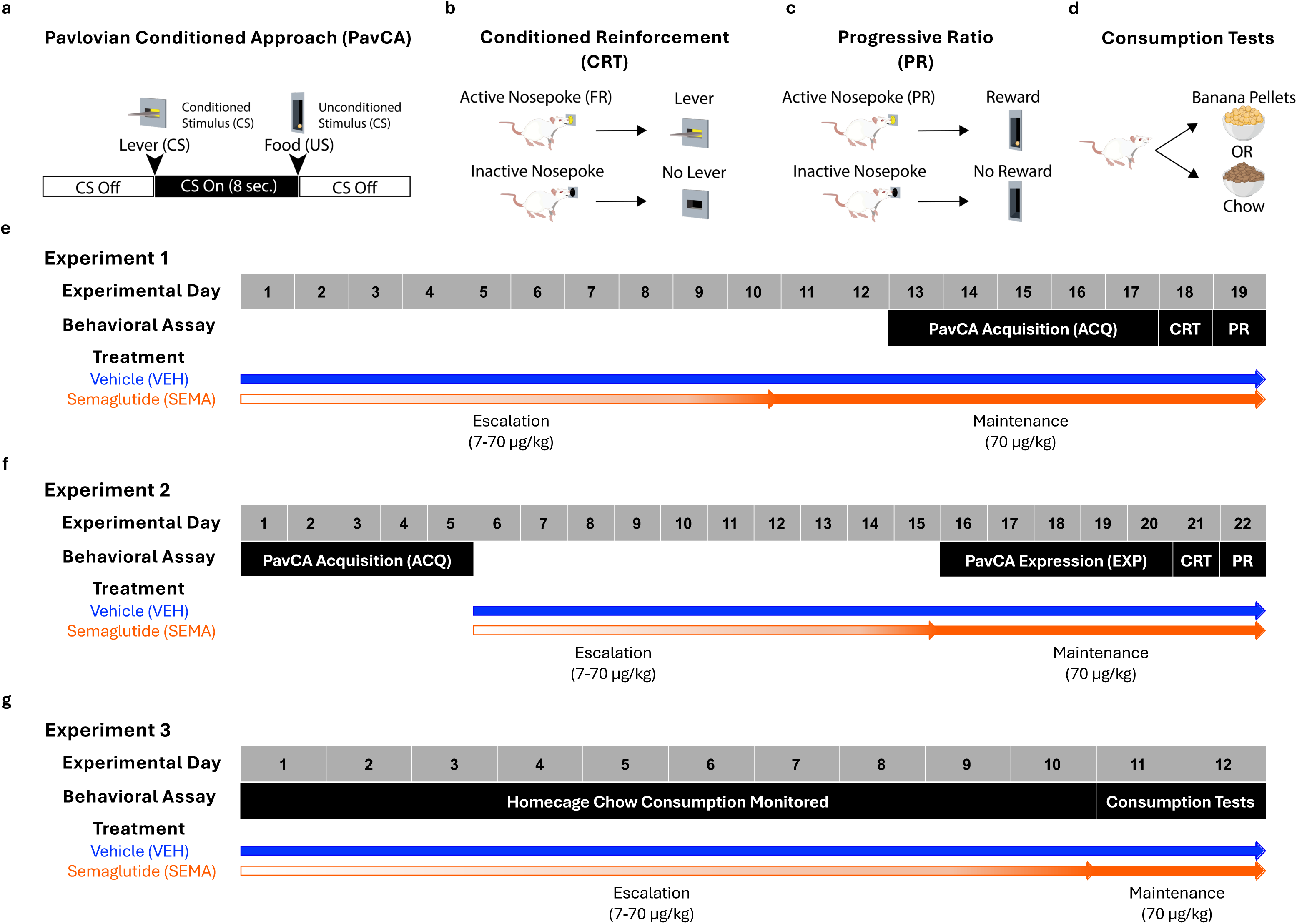
Experimental timeline. Schematics showing (a) Pavlovian Conditioned Approach (PavCA), (b) Conditioned Reinforcement (CRT), (c) Progressive Ratio (PR) Responding, and (d) Consumption Tests. (e) Experiment 1 investigated the effects of chronic semaglutide on the *acquisition* of PavCA behavior, CRT, and PR responding. (f) Experiment 2 investigated the effects of chronic semaglutide on the *expression* of PavCA behavior, CRT, and PR responding. (g) Experiment 3 investigated the effects of chronic semaglutide on the free consumption of chow and banana-flavored pellets used in Experiments 1 and 2.

### Experiment 1

The purpose of Experiment 1 was to assess the effects of chronic semaglutide administration on the *acquisition* of PavCA behavior, the conditioned reinforcing properties of the food-associated cue, and the reinforcing properties of the food reward. The experimental timeline is illustrated in **Figure 1e**. Behavioral testing occurred between the hours of 11:00 AM and 1:00 PM, during the rats’ light phase.

#### Subjects

The subjects were adult Sprague Dawley rats (n = 64; 32 male and 32 female) from Charles River (Stone Ridge, NY). The males were from Barriers K90 and K98 (16 each), while all female rats were from Barrier K90. Experiment 1 was conducted by SC and SK. Rats were run in 2 rounds of 32 (16 males and 16 females each round), spaced 1 month apart. Rats weighed 275-325g on arrival and were given *ad libitum* access to food and water for the duration of the experiment. Rats were pair-housed and allowed to acclimate to the climate-controlled vivarium (12-h light:dark cycle with lights on at 7:00 AM and lights off at 7:00 PM) for 1 week prior to experimental procedures.

#### Semaglutide Onboarding

The chronic semaglutide treatment regimen was adapted from Cawthon et al. (2023). Rats received daily subcutaneous injections of either vehicle (VEH; 44mM sodium phosphate dibasic, 70 mM NaCl + 0.007% Tween 20) or semaglutide (SEMA; Astatech) diluted in the vehicle solution. Half the rats were assigned to the VEH group (16 male, 16 female) and the other half were assigned to the SEMA group (16 male, 16 female). Injections occurred at approximately 1:00 PM each day, after behavioral assays to minimize the impact of acute effects on behavior. Semaglutide administration started at a dosage of 7 µg/kg body weight and escalated 7 µg/kg each day until reaching the final dose of 70 µg/kg body weight (10-day period). Rats then remained on a 70 µg/kg body weight dose per day for the remainder of the experiment. Behavioral testing started once rats reached the 70 µg/kg dose. Body weights were recorded each day, and the percent change in body weight from baseline (i.e., Day 0 weight) was calculated for the duration of the experiment.

#### Pavlovian Conditioned Approach (PavCA)

Methods for PavCA training are similar to those previously described (Robinson & Flagel, 2009). Prior to behavioral testing, rats were given 45-mg grain-based banana-flavored pellets in their home cages (approximately 50 pellets per cage; Bioserv; Product #F0059) for 2 days, to familiarize them with the food reward. PavCA training occurred in standard behavioral chambers from Med Associates (20.5 cm x 24.1 cm x 29.2 cm) that were encased in sound-attenuating boxes with ventilation fans that provided air circulation and background noise. A recessed food cup was located in the center of one wall 1.5 cm above the grid floor. A retractable lever-cue was located to the left or right of the food cup 6 cm above the grid floor (counterbalanced to control for side bias). The lever-cue was illuminated with an LED backlight upon presentation. A houselight was located on the opposite wall 1 cm from the top of the chamber and was illuminated throughout each session.

Rats were given 2 pre-training sessions in which one banana-flavored reward pellet was delivered randomly into the food cup on a variable interval 30-s schedule (range 0 – 60 s; 25 pellets total per session). Following pre-training, rats underwent 5 sessions of PavCA training. Each session consisted of 25 trials in which the retractable illuminated lever-cue (conditioned stimulus, CS) was presented for 8 s and, upon retraction, a food reward pellet (unconditioned stimulus, US) was immediately delivered. Trials were presented on a variable interval 90-s schedule (range 30 – 150 s).

The following measures were collected via Med Associates software for each session: total number of lever-cue contacts, total number of food cup entries during lever-cue presentations, latency to the first lever-cue contact, and latency to the first food cup entry during lever-cue presentation. These measures allowed us to calculate the PavCA Score (Meyer et al., 2012) for each rat based on the: Response Bias [(lever contacts – food cup entries)/(lever contacts + food cup entries)], Probability Difference (p|lever contact] – p|food cup entry]), and Latency Score [(average response time to lever contact – average response time to enter the food cup)/duration of the lever-cue presentation]. The average of these three measures was calculated to determine the PavCA Score for each of the 5 sessions, which ranged from -1.0 to 1.0. A score of +1 indicates that a rat is solely interacting with the lever-cue upon its presentation (i.e., sign-tracking), and a score of -1 indicates that a rat is solely interacting with the food cup upon lever-cue presentation (i.e., goal-tracking). A score around 0 suggests that a rat does not have a preference for either the lever-cue or the food cup. The PavCA Index is then calculated as the average PavCA Score from Sessions 4 and 5, after rats have acquired their conditioned response (Meyer et al., 2012).

#### Conditioned Reinforcement Test (CRT)

The day following PavCA training, rats underwent a conditioned reinforcement test (CRT) to assess the reinforcing properties of the lever-cue, as previously described (Robinson & Flagel, 2009). The food cup was removed from the center wall and replaced with the retractable lever-cue used during PavCA training. Nose ports were placed to the left and right of the lever-cue (one on each side; 6 cm above the grid floor). One port was designated as the active port and the other was designated as the inactive port. The active port was placed on the opposite side of where the lever-cue was located during PavCA training to avoid side bias. During the 45-min conditioned reinforcement test, pokes into the active nose port resulted in a 2-sec presentation of the retractable lever that was previously associated with food reward. Pokes into the inactive nose port had no consequence.

The following measures were collected using Med Associates software: the total number of nose pokes into the active and inactive ports, and total number of lever contacts upon its presentation. These measures were then used to calculate the Incentive Value Index [(pokes into the active port + lever contacts) – pokes into the inactive port], as previously described (Hughson et al., 2019).

#### Progressive Ratio (PR) Test

The 32 rats (16 male, 16 female) from round 2 of Experiment 1 underwent testing for progressive ratio (PR) responding to assess motivation for the banana-flavored reward pellet. The day following the conditioned reinforcement test, the retractable lever was removed from the chambers and the food cup was put back in the center of the wall (6 cm above the grid floor) with nose ports on either side (6 cm above the grid floor). Pokes into the active port resulted in delivery of a single reward pellet, while pokes into the inactive port had no consequence. Active and inactive port assignments were the same as during the conditioned reinforcement test. Pellet delivery was contingent upon pokes into the active port based on the progressive ratio schedule: 1,1,1,2,2,2,3,3,3,4,4,4,5,5,5,10,15,20,25,30,35,40,45,50,55,60,65,70,75,80,85,90,95,100,110,12 0,130,140,150,160,170,180,190,200,220,240,260,280,300,330,360,390,420,460,500,550,600,6 60,720,790,860,940,1030,2030. The following measures were collected: total number of nose pokes in the active and inactive ports, total number of rewards received, and breakpoint, defined as the last PR reached before session termination. The session terminated if 30 min elapsed without meeting a PR or if the total session length reached 90 min.

#### Data Analysis

For Experiment 1, the percent change in body weight from baseline and PavCA data were analyzed using linear mixed models (LMMs) with factors of Treatment (VEH vs. SEMA), Sex (Male vs. Female), and Session (Percent Weight Change: 1-17; PavCA: 1-5). Subject was included as a random intercept. These analyses were conducted using SPSS Statistics, Version 31 (IBM, Armonk, NY). PavCA Index data (average of PavCA Scores of Sessions 4 and 5) were analyzed using ANOVAs with factors of Treatment (VEH vs. SEMA) and Sex (Male vs. Female). Conditioned reinforcement nosepoke data were analyzed using ANOVAs with factors of Port (Active vs. Inactive), Treatment (VEH vs. SEMA), and Sex (Male vs. Female). Lever contacts and the Incentive Value Index were analyzed using ANOVAs with factors of Treatment (VEH vs. SEMA) and Sex (Male vs. Female). Progressive ratio nosepoke data was analyzed using ANOVAs with the factors of Port (Active vs. Inactive), Treatment (VEH vs. SEMA), and Sex (Male vs. Female). Reward total, breakpoint data, and were analyzed using ANOVAs with factors of Treatment (VEH vs. SEMA) and Sex (Male vs. Female). Any statistically significant main effects and interactions were followed up with *post hoc* comparisons (t-tests) with Bonferroni corrections as appropriate. Effect sizes for pairwise comparisons were calculated using Cohen’s d (Cohen, 1988), as previously reported (Hughson et al., 2019). Effect sizes < 0.2 are considered small, effect sizes ≥ 0.2 and ≤ 0.8 are considered medium, and effect sizes > 0.8 are considered large (Cohen, 1988; Sawilowsky, 2009). Pairwise Pearson correlation coefficients were calculated using GraphPad Prism, Version 10.6.1 (GraphPad Software, San Diego, CA) to examine potential linear relationships between responding during the conditioned reinforcement test and the progressive ratio test. Data for Experiment 1 are shown in **Figure 2** and **Table 1**.

**Figure 2.**
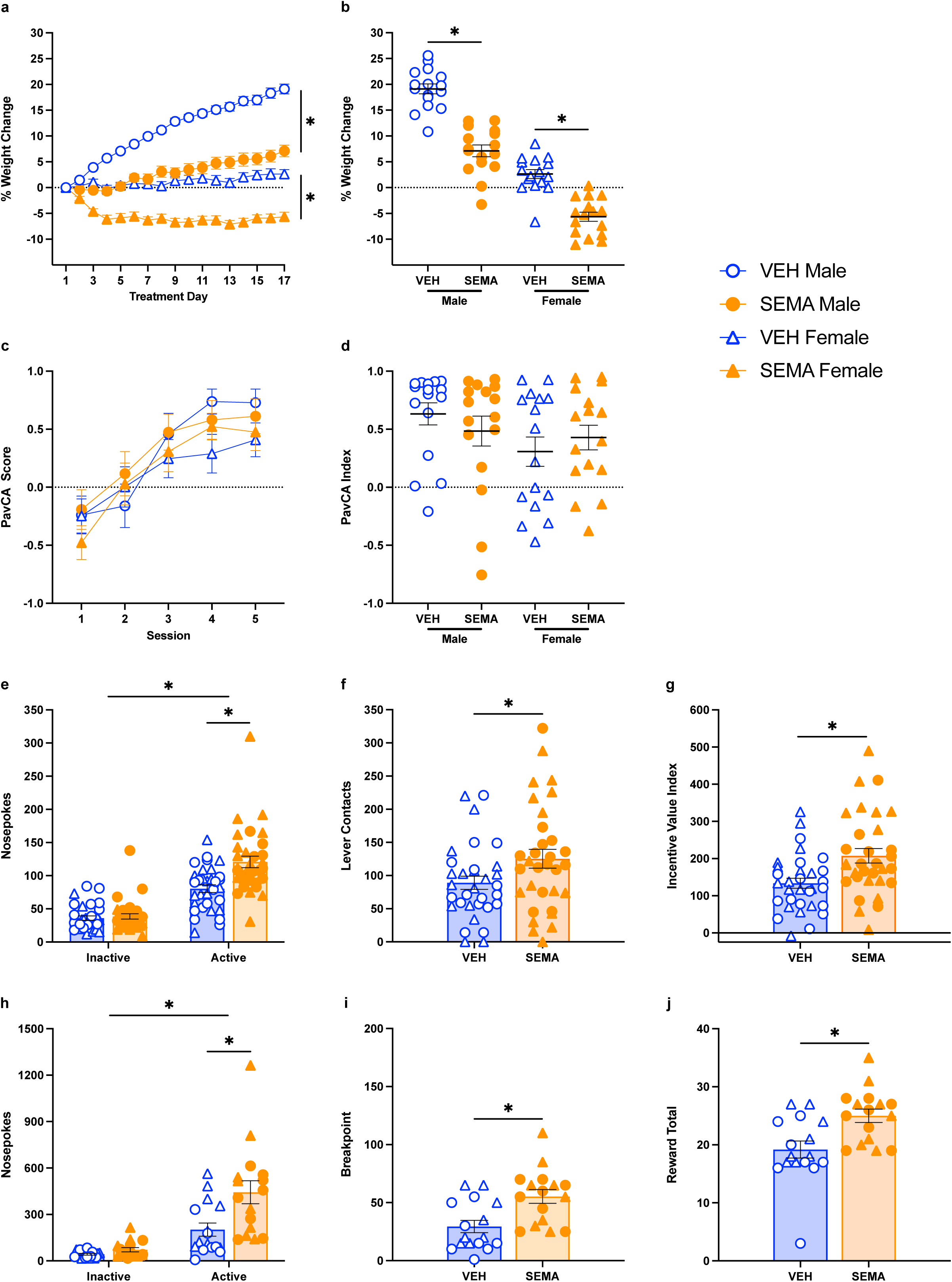
The effects of chronic semaglutide treatment on (a-b) body weight, (c-d) the *acquisition* of Pavlovian conditioned approach behavior, (e-g) the conditioned reinforcing properties of a food-paired cue, and (h-j) progressive ratio responding for food reward. (a-g) Data are shown for vehicle- (VEH, n = 32; 16 males and 16 females) and semaglutide- (SEMA, n = 32; 16 males and 16 females) treated rats. (a) Mean ± SEM percent change in weight from baseline for VEH- and SEMA-treated rats across 17 days of treatment. Relative to vehicle, chronic semaglutide treatment reduced weight gain in males and induced weight loss in females. (b,d) Black bars show mean ± SEM and (b,d-j) each data point represents an individual rat. (b) Weight change from baseline following the final treatment on Day 17. Semaglutide treatment reduced weight gain relative to vehicle-treated rats for both males and females. (c) Mean ± SEM PavCA Scores of VEH- and SEMA-treated rats over 5 PavCA Sessions. Chronic semaglutide treatment had no effect on the *acquisition* of PavCA behavior. (d) Data show PavCA Index, the average of PavCA Scores from Sessions 4 and 5. There were no significant differences in PavCA Index between groups. (e) Mean ± SEM nosepokes into the active and inactive ports for VEH- and SEMA-treated rats during conditioned reinforcement. Both groups responded more in the active port relative to the inactive port. Chronic semaglutide treatment enhanced responding in the active nose port relative to vehicle. (f) Mean ± SEM lever contacts during conditioned reinforcement. SEMA-treated rats made more lever contacts than VEH-treated rats. (g) Mean ± SEM incentive value index during conditioned reinforcement. SEMA-treated rats had higher incentive value indexes than VEH-treated rats. (h-j) Data are shown for a subset of 32 rats (n=16 per group, n=8 per sex) that were assessed for progressive ratio responding. Mean ± SEM nosepokes into the active and inactive ports of VEH-and SEMA-treated rats during progressive ratio responding. Both groups responded more in the active port relative to the inactive port, and SEMA-treated rats responded more in the active nose port relative to VEH-treated rats. (i) Mean ± SEM breakpoint of VEH- and SEMA-treated rats during progressive ratio responding. SEMA-treated rats had higher breakpoints than VEH-treated rats. (j) Mean ± SEM total rewards received for VEH- and SEMA-treated rats during the progressive ratio test. SEMA-treated rats received more rewards than VEH-treated rats. \**p* < 0.05

**Table 1.** Statistical analyses for (a) lever-cue- and (b) food-cup-directed behaviors for Experiment 1. Linear mixed models for sessions 1-5 of PavCA (degrees of freedom (DF), F-value, P-value). Significant effects (*p* < 0.05) are indicated in bold with an asterisk.

| a | Lever-cue Contacts |  |  | Probability Lever-cue Contact |  |  | Latency Lever-cue Contact |  |  |
| --- | --- | --- | --- | --- | --- | --- | --- | --- | --- |
|  | DF | F | P | DF | F | P | DF | F | P |
| Sex | 1,47 | 0.837 | 0.365 | 1,62 | 0.682 | 0.412 | 1,60 | 1.132 | 0.292 |
| Treatment | 1,47 | 0.105 | 0.747 | 1,62 | 0.123 | 0.727 | 1,60 | 0.268 | 0.607 |
| Session | 4.131 | 30.349 | **\*<0.001** | 4,141 | 41.493 | **\*<0.001** | 4,146 | 41.085 | **\*<0.001** |
| Treatment * Session | 4.131 | 0.535 | 0.711 | 4,141 | 0.508 | 0.73 | 4,146 | 0.461 | 0.764 |
| Sex * Session | 4.131 | 1.692 | 0.156 | 4,141 | 1.014 | 0.402 | 4,146 | 1.36 | 0.251 |
| Sex * Treatment | 1,47 | 0.232 | 0.632 | 1,62 | 0.069 | 0.793 | 1,60 | 0.03 | 0.864 |
| Sex * Treatment * Session | 4.131 | 0.978 | 0.422 | 4,141 | 2.281 | 0.064 | 4,146 | 1.477 | 0.212 |
| b | Food-cup Entry |  |  | Probability Food-cup Entry |  |  | Latency Food-cup Entry |  |  |
|  | DF | F | P | DF | F | P | DF | F | P |
| Sex | 1,64 | 0.09 | 0.765 | 1,61 | 1.012 | 0.318 | 1,78 | 0.78 | 0.38 |
| Treatment | 1,64 | 1.359 | 0.248 | 1,61 | 0.622 | 0.433 | 1,78 | 1.402 | 0.24 |
| Session | 4,122 | 7.993 | **\*<0.001** | 4,73 | 6.856 | **\*<0.001** | 4,94 | 8.289 | **\*<0.001** |
| Treatment * Session | 4,122 | 0.523 | 0.719 | 4,73 | 0.221 | 0.926 | 4,94 | 0.29 | 0.884 |
| Sex * Session | 4,122 | 0.384 | 0.82 | 4,73 | 1.027 | 0.399 | 4,94 | 0.544 | 0.704 |
| Sex * Treatment | 1,64 | 0.805 | 0.373 | 1,61 | 1.478 | 0.229 | 1,78 | 0.81 | 0.371 |
| Sex * Treatment * Session | 4,122 | 1.745 | 0.144 | 4,73 | 1.879 | 0.123 | 4,94 | 2.502 | **\*0.048** |

### Experiment 2

The purpose of Experiment 2 was to assess the effects of chronic semaglutide administration on the *expression* of PavCA behavior, the conditioned reinforcing properties of the food-associated cue, and the reinforcing properties of the food reward itself. The experimental timeline is illustrated in **Figure 1f**. Behavioral testing occurred between the hours of 11:00 AM and 1:00 PM, during the rats’ light cycle.

#### Subjects

The subjects were adult Sprague Dawley rats (n = 64; 32 male and 32 female) from Charles River (16 males, 16 females from Barriers K98 in Stone Ridge, NY and 16 males, 16 females from Barrier R09 in Raleigh, NC). Experiment 2 was conducted by SC, NM, DP, and CT. Rats were run in two rounds (with equal representation of both sexes from both Barriers) spaced 2 months apart. Rats weighed 275-325g on arrival and were given *ad libitum* access to food and water for the duration of the experiment. Rats were pair-housed and allowed to acclimate to the climate-controlled vivarium (12-h light:dark cycle with lights on at 7:00 AM and lights off at 7:00 PM) for 1 week prior to experimental procedures. One male rat was excluded prior to the start of vehicle/semaglutide injections, due to not consuming the banana-flavored reward pellets during PavCA training.

#### Pavlovian Conditioned Approach (PavCA): Acquisition Phase

For Experiment 2, rats underwent PavCA training before chronic semaglutide administration commenced. As described for Experiment 1, rats were pre-exposed to the banana-flavored reward pellets for 2 days (approximately 50 pellets per cage) prior to behavioral testing. Rats then underwent pre-training and 5 Sessions of PavCA training as in Experiment 1. Assessing PavCA behavior prior to chronic semaglutide administration allowed us to classify rats based on their behavioral phenotypes (i.e., whether rats were sign-trackers or goal-trackers). This was done through calculating the PavCA Index, which is the average of PavCA Scores from Sessions 4 and 5 (Meyer et al., 2012). Rats with a PavCA Index greater than +0.5 were classified as sign-trackers, and rats with a PavCA Index less than -0.5 were classified as goal-trackers. Rats with a PavCA Index between -0.5 and +0.5 were classified as intermediate responders.

#### Semaglutide Onboarding

Following PavCA training, chronic semaglutide administration began using the same regimen as in Experiment 1. Half of the rats were assigned to the VEH group while the other half were assigned to the SEMA group. Group assignments were counterbalanced based on the PavCA Index to ensure that each group had a similar distribution in terms of behavioral phenotypes when PavCA training recommenced. This resulted in 31 vehicle- (16 males, 15 females) and 32 semaglutide- (15 males, 17 females) treated rats.

#### Pavlovian Conditioned Approach (PavCA): Expression Phase

Once rats completed semaglutide onboarding across 10 days, they underwent 5 more Sessions of PavCA training while receiving the highest dose of semaglutide (70 µg/kg) or VEH injections. As in Experiment 1, injections occurred daily, after behavioral testing, around 1:00 PM.

#### Conditioned Reinforcement Test

The day following the last PavCA session, all rats underwent testing for CRT as described in Experiment 1, after receiving semaglutide (70 µg/kg) or VEH injections the day prior to the test.

#### Progressive Ratio Test

The day following CRT, all rats underwent testing for PR responding as described in Experiment 1, after receiving semaglutide (70 µg/kg) or VEH injections the day prior to the test.

#### Data Analysis

For Experiment 2, percent change in body weight from baseline and PavCA data were analyzed using LMMs with factors of Treatment (VEH vs. SEMA), Sex (Male vs. Female), and Treatment Day (Percent Weight Change: 1-17) or Session (PavCA: 1-5 and 6-10). Subject was included as a random intercept. These analyses were conducted using SPSS Statistics, Version 31 (IBM, Armonk, NY). PavCA Index data (average of PavCA Scores of Sessions 4 and 5 as well as Sessions 9 and 10) were analyzed using ANOVAs with factors of Treatment (VEH vs. SEMA) and Sex (Male vs. Female). Conditioned reinforcement and progressive ratio data were analyzed as described for Experiment 1. The significance level was *p* < 0.05 for all tests. Any statistically significant main effects and interactions were followed up with *post hoc* comparisons (t-tests) with Bonferroni corrections as appropriate. Effect sizes for pairwise comparisons were calculated using Cohen’s d (Cohen, 1988), as previously reported (Hughson et al., 2019). Pairwise Pearson correlation coefficients were calculated using GraphPad Prism, Version 10.6.1 (GraphPad Software, San Diego, CA) to examine potential linear relationships between responding during the conditioned reinforcement test and the progressive ratio test. Data for Experiment 2 are shown in **Figure 3** and **Table 2**.

**Figure 3.**
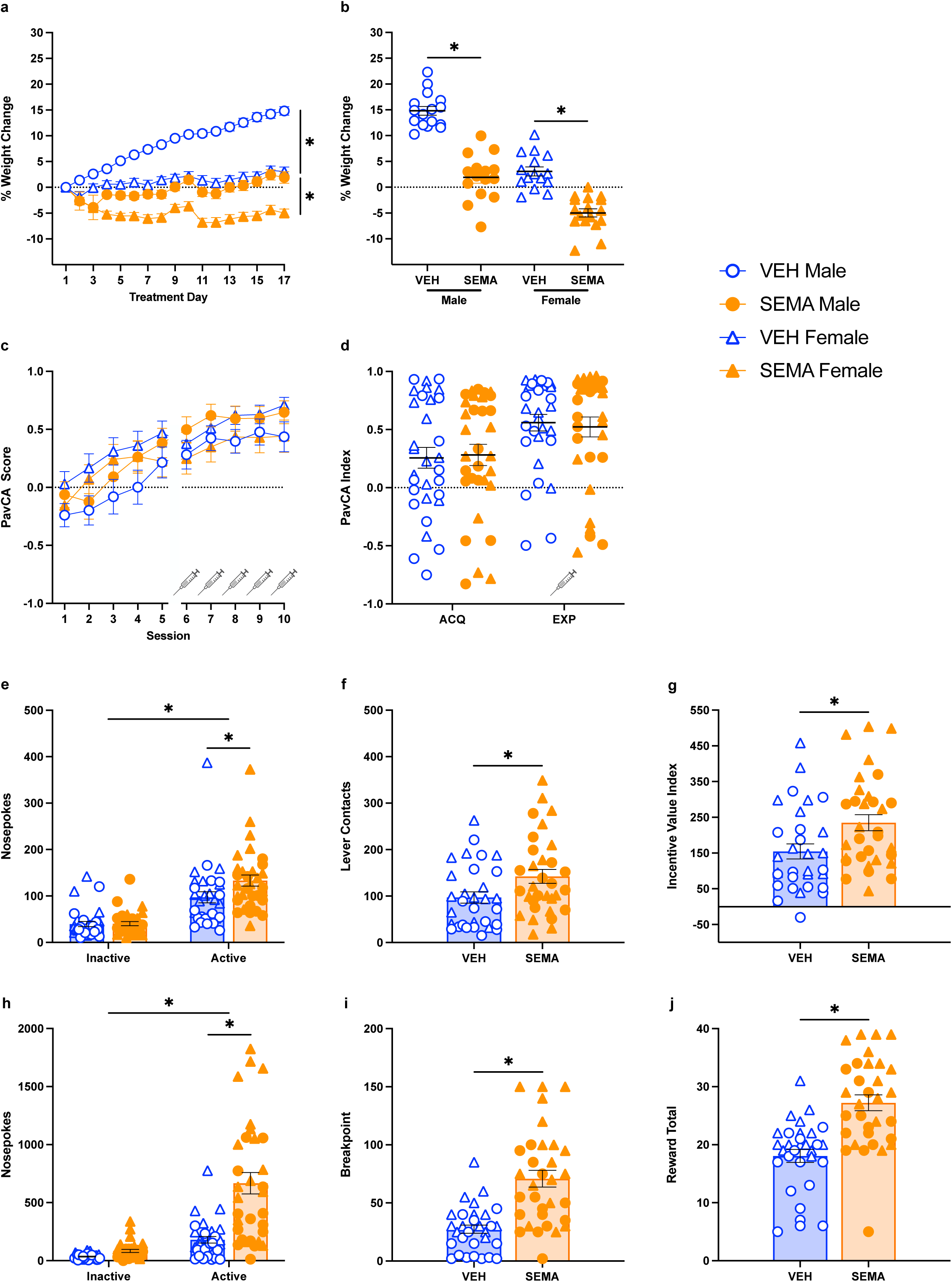
The effects of chronic semaglutide treatment on (a-b) body weight, (c-d) the expression of Pavlovian conditioned approach behavior, (e-g) the conditioned reinforcing properties of a food-paired cue, and (h-j) progressive ratio responding for food reward. (a-g) Data are shown for vehicle- (VEH, n = 31; 16 males and 15 females) and semaglutide- (SEMA, n = 32; 15 males and 17 females) treated rats. (a) Mean ± SEM percent change in weight from baseline for VEH- and SEMA-treated rats across 17 days of treatment. Relative to vehicle, chronic semaglutide treatment reduced weight gain in males and induced weight loss in females relative to vehicle. (b,d) Black bars show mean ± SEM and (b,d-j) each data point represents an individual rat. (b) Weight change from baseline following the final treatment on Day 17. Semaglutide treatment reduced weight gain relative to vehicle-treated rats for both males and females. (c) Mean ± SEM PavCA Scores of VEH- and SEMA-treated rats over PavCA Sessions 1-5 prior to treatment, and 6-10 during treatment (indicated with syringe). Chronic semaglutide treatment had no effect on the *expression* of PavCA behavior over sessions 6-10. (d) Data show PavCA Index, the average of PavCA Scores from Sessions 4 and 5 during acquisition (ACQ) and the average from Sessions 9 and 10 during expression (EXP) There were no significant differences in PavCA Index between groups or phases (i.e., ACQ vs EXP). (e) Mean ± SEM nosepokes into the active and inactive ports for VEH- and SEMA-treated rats during conditioned reinforcement. Both groups responded more in the active port relative to the inactive port. Chronic semaglutide treatment enhanced responding in the active nose port relative to vehicle. (f) Mean ± SEM lever contacts during conditioned reinforcement. SEMA-treated rats made more lever contacts than VEH-treated rats. (g) Mean ± SEM incentive value index during conditioned reinforcement. SEMA-treated rats had higher incentive value indexes than VEH rats. (h) Mean ± SEM nosepokes into the active and inactive ports of VEH and SEMA rats during progressive ratio responding. Both groups responded more in the active port relative to the inactive port., and SEMA-treated rats responded more in the active nose port relative to VEH-treated rats. (i) Mean ± SEM breakpoint for VEH- and SEMA-treated rats during the progressive ratio test. SEMA-treated rats had higher breakpoints than VEH-treated rats. (j) Mean ± SEM total rewards received for VEH- and SEMA-treated rats during the progressive ratio test. SEMA-trated rats received more rewards than VEH-treated rats. \**p* < 0.05

**Table 2.** Statistical analyses for (a) lever-cue- and (b) food-cup-directed behaviors for Experiment 2. Linear mixed models for sessions 1-5 and sessions 6-10 of PavCA (degrees of freedom (DF), F-value, P-value). Significant effects (*p* < 0.05) are indicated in bold with an asterisk.

**Sessions 1-5: Acquisition**
| **a** | Lever-cue Contacts |  |  | Probability Lever-cue Contact |  |  | Latency Lever-cue Contact |  |  |
| --- | --- | --- | --- | --- | --- | --- | --- | --- | --- |
|  | DF | F | P | DF | F | P | DF | F | P |
| Sex | 1,59 | 3.269 | 0.076 | 1,64 | 6.083 | **\*0.016** | 1,67 | 3.987 | 0.5 |
| Treatment | 1,59 | 0.126 | 0.723 | 1,64 | 0.007 | 0.934 | 1,67 | 0.044 | 0.834 |
| Session | 4,108 | 19.822 | **\*<0.001** | 4,91 | 23.846 | **\*<0.001** | 4,85 | 19.197 | **\*<0.001** |
| Treatment *<br>Day | 4,108 | 0.294 | 0.881 | 4,91 | 0.575 | 0.681 | 4,85 | 0.651 | 0.628 |
| Sex *<br>Session | 4,108 | 1.608 | 0.178 | 4,91 | 1.631 | 0.173 | 4,85 | 1.114 | 0.355 |
| Sex *<br>Treatment | 1,59 | 2.272 | 0.137 | 1,63 | 2.769 | 0.101 | 1,67 | 1.451 | 0.233 |
| Sex *<br>Treatment *<br>Session | 4,108 | 0.953 | 0.436 | 4,91 | 1.44 | 0.227 | 4,85 | 0.802 | 0.527 |

| **b** | Food-cup Entry |  |  | Probability Food-cup Entry |  |  | Latency Food-cup Entry |  |  |
| --- | --- | --- | --- | --- | --- | --- | --- | --- | --- |
|  | DF | F | P | DF | F | P | DF | F | P |
| Sex | 1,60 | 1.201 | 0.278 | 1,75 | 0.009 | 0.927 | 1,74 | 0.009 | 0.924 |
| Treatment | 1,60 | 0.019 | 0.89 | 1,75 | 0.08 | 0.778 | 1,74 | 0.002 | 0.966 |
| Session | 4,72 | 2.286 | 0.068 | 4,87 | 4.212 | **\*0.004** | 4,87 | 3.399 | **\*0.012** |
| Treatment *<br>Day | 4,72 | 1.815 | 0.135 | 4,87 | 1.11 | 0.357 | 4,87 | 1.765 | 0.143 |
| Sex *<br>Session | 4,72 | 1.506 | 0.209 | 4,87 | 1.144 | 0.342 | 4,87 | 0.813 | 0.52 |
| Sex *<br>Treatment | 1,60 | 0.709 | 0.403 | 1,75 | 0.938 | 0.336 | 1,74 | 0.524 | 0.471 |
| Sex *<br>Treatment *<br>Session | 4,72 | 1.5 | 0.211 | 4,87 | 0.325 | 0.861 | 4,87 | 0.414 | 0.798 |

**Sessions 6-10:** Expression
| **a** | Lever-cue Contacts |  |  | Probability Lever-cue Contact |  |  | Latency Lever-cue Contact |  |  |
| --- | --- | --- | --- | --- | --- | --- | --- | --- | --- |
|  | DF | F | P | DF | F | P | DF | F | P |
| Sex | 1,61 | 0.47 | 0.495 | 1,58 | 0.216 | 0.644 | 1,60 | 0.001 | 0.977 |
| Treatment | 1,61 | 1.809 | 0.184 | 1,58 | 0.235 | 0.63 | 1,60 | 0.622 | 0.433 |
| Session | 4,83 | 3.865 | **\*0.006** | 4,114 | 3.718 | **\*0.007** | 4,73 | 5.038 | **\*0.001** |
| Treatment *<br>Day | 4,83 | 0.171 | 0.953 | 4,114 | 0.111 | 0.978 | 4,73 | 0.219 | 0.927 |
| Sex *<br>Session | 4,83 | 2.894 | **\*0.027** | 4,114 | 3.016 | **\*0.021** | 4,73 | 2.8 | **\*0.032** |
| Sex *<br>Treatment | 1,61 | 4.154 | **\*0.046** | 1,58 | 2.959 | 0.091 | 1,60 | 2.218 | 0.142 |
| Sex *<br>Treatment *<br>Session | 4,83 | 0.982 | 0.422 | 4,114 | 2.637 | **\*0.038** | 4,73 | 1.963 | 0.109 |

| **b** | Food-cup Entry |  |  | Probability Food-cup Entry |  |  | Latency Food-cup Entry |  |  |
| --- | --- | --- | --- | --- | --- | --- | --- | --- | --- |
|  | DF | F | P | DF | F | P | DF | F | P |
| Sex | 1,59 | 0.069 | 0.793 | 1,62 | 0.006 | 0.939 | 1,62 | 0.005 | 0.944 |
| Treatment | 1,59 | 0.205 | 0.652 | 1,62 | 0.014 | 0.905 | 1,62 | 0.003 | 0.953 |
| Session | 4,32 | 6.034 | **\*<0.001** | 4,139 | 5.71 | **\*<0.001** | 4,134 | 9.026 | **\*<0.001** |
| Treatment *<br>Day | 4,32 | 0.238 | 0.915 | 4,139 | 0.584 | 0.675 | 4,134 | 0.264 | 0.9 |
| Sex *<br>Session | 4,32 | 0.5 | 0.736 | 4,139 | 1.087 | 0.366 | 4,134 | 0.933 | 0.447 |
| Sex *<br>Treatment | 1,59 | 1.593 | 0.212 | 1,62 | 1.495 | 0.226 | 1,62 | 2.212 | 0.142 |
| Sex *<br>Treatment *<br>Session | 4,32 | 0.432 | 0.785 | 4,139 | 0.612 | 0.655 | 4,134 | 0.944 | 0.441 |

### Experiment 3

The purpose of Experiment 3 was to assess the effects of chronic semaglutide administration on homecage chow consumption and free consumption of chow or banana-flavored reward pellets. The experimental timeline is illustrated in **Figure 1g**.

#### Subjects

The subjects were adult Sprague Dawley rats (n = 32; 16 male and 16 female) from Charles River (Barriers K98 in Stone Ridge, NY and R06 in Raleigh, NC). The males were from Barrier K98, and the females were from Barrier R06. Experiment 3 was conducted in one round by SC. Rats weighed 275-325g on arrival and were given *ad libitum* access to food and water for the duration of the experiment. Rats were pair-housed and allowed to acclimate to the climate-controlled vivarium (12-h light:dark cycle with lights on at 7:00 AM and lights off at 7:00 PM) for 1 week prior to experimental procedures.

#### Semaglutide Onboarding and Homecage Consumption

Chronic semaglutide or VEH administration occurred as described above for Experiments 1 and 2, around 1:00 PM each day. In addition to body weight, homecage food intake was recorded each day, immediately prior to injections. To assess food intake for each group, rats in the same cage were assigned to the same treatment group (VEH or SEMA). During the last 7 days of semaglutide onboarding (i.e., doses 28-70 µg/kg) rats received banana-flavored reward pellets (approximately 50 per cage) in their homecage to simulate the amount of reward pellets received during Experiments 1 and 2.

#### Free-feeding Consumption Tests

The day following completion of the semaglutide onboarding regimen, rats underwent two free-feeding consumption tests. Rats were first habituated to the unfamiliar test environment, which was an acrylic cage (43 x 21.5 x 25.5 cm (high) with a metal grid floor insert and cage top. Cages were on racks separated by black acrylic walls to minimize visual contact between subjects. An empty ceramic bowl was placed in the front right corner of each cage. For three consecutive days, rats were placed in these cages at 12:00 PM for a 30-min session to allow for habituation to the testing environment.

The day following the last habituation session, food was removed from the homecages at 8:00 AM, to control for the amount of food eaten and motivational state of the rats immediately prior to the consumption tests. On the first test day, at 12:00 PM, rats were placed into the test chambers and given *ad libitum* access to either banana-flavored reward pellets or chow for 15 min. Half of the rats received banana-flavored pellets on the first session, while the other half received chow. The food exposure was reversed for the second session. The amount of food in the ceramic bowls was weighed before and after each session to calculate the amount of food consumed.

#### Data Analysis

For Experiment 3, percent change in body weight from baseline data was analyzed using LMMs with factors of Treatment (VEH vs. SEMA), Sex (Male vs. Female), and Day (1-15). The amount of food consumed during semaglutide treatment was averaged into 2-Day bins and analyzed per cage using LMMs with factors of Treatment (VEH vs. SEMA), Sex (Male vs. Female), and Bin (1-7). Subject was included as a random intercept for all LMMs. These analyses were conducted using SPSS Statistics, Version 31 (IBM, Armonk, NY). For the consumption tests, the amount of food consumed was analyzed using ANOVAs with factors of Treatment (VEH vs. SEMA), Sex (Male vs. Female), and Food Type (Chow vs. Banana Pellets). The significance level was *p* < 0.05 for all tests. Any statistically significant main effects and interactions were followed up with *post hoc* comparisons (t-tests) with Bonferroni corrections as appropriate. Effect sizes for pairwise comparisons were calculated using Cohen’s d (Cohen, 1988), as previously reported (Hughson et al., 2019).

## Results

### Experiment 1

#### Weight Change

Relative to vehicle-treated rats, chronic semaglutide treatment reduced weight gain in males and induced weight loss in females (**Figure 2a-b**). This is supported by significant main effects of Treatment (F_1,60_ = 130.24, *p* < 0.001), Day (F_16,960_ = 79.41, *p* < 0.001), and Sex (F_1,60_ = 196.85, *p* < 0.001), as well as significant Treatment x Day (F_16,960_ = 41.41, *p* < 0.001) and Day x Sex (F_16,960_ = 97.30, *p* < 0.001) interactions. After Day 1 of treatment, SEMA-treated rats (regardless of sex) weighed significantly less than VEH-treated rats for the remainder of the experiment (*p* ≤ 0.006) (**Figure 2a**). At the end of the 17-day treatment regimen, both male and female rats treated with semaglutide had reduced weight gain with respect to baseline body weight relative to those treated with vehicle (**Figure 2b**; Males: t = 8.038, *p* < 0.001; Cohen’s d = 2.842; Females: t = 6.668, *p* < 0.001; Cohen’s d = 2.358). There was a significant effect of Treatment (F_1,60_ = 109.1, *p* < 0.001) and Sex (F_1,60_ = 226.9, *p* < 0.001), but no Treatment x Sex interaction (F_1,60_ = 3.7, *p* = 0.059) for body weight on the final day of treatment. As evident in **Figure 2b**, most female rats treated with semaglutide showed a decrease in body weight relative to baseline levels, whereas most male rats showed attenuated weight gain rather than weight loss. For both sexes, however, there was approximately a 10% difference in the change from baseline body weight in rats treated with semaglutide relative to those treated with vehicle, demonstrating the validity of the treatment regimen.

#### Pavlovian Conditioned Approach (PavCA)

Chronic semaglutide administration had no effect on the acquisition of PavCA behavior (**Figure 2c**). Table 1 presents the LMM analyses of the individual behavioral measures (i.e., lever-cue contacts, probability of lever-cue contact, latency to lever-cue contact, food-cup entries, probability of food-cup entry, and latency to food-cup entry) used to calculate PavCA Scores. For each measure, there was a significant main effect of Session (*p* < 0.001), but no main effects of Sex (*p* ≥ 0.292) or Treatment (*p* ≥ 0.240).

As shown in **Figures 2c-d**, by the end of PavCA training, all groups tend to sign-track (i.e., PavCA Score > 0.5) rather than goal-track. There was a significant effect of Session (F_4,88_ = 29.82, *p* < 0.001), but no significant main effects of Treatment (F_1,63_ = 0.01, *p* = 0.943) or Sex (F_1,63_ = 1.53, *p* = 0.220) for the PavCA Score (**Figure 2c**). The PavCA Index (i.e., average of PavCA Score on Sessions 4 and 5) also did not differ between treatment groups or sex (Treatment (F_1,60_ = 0.01, *p* = 0.907); Sex (F_1,60_ = 2.75, *p* = 0.102)), and there were no significant interactions (Treatment x Sex interaction (F_1,60_ = 1.38, *p* = 0.246)) (**Figure 2d**). Thus, chronic semaglutide treatment does not appear to affect the acquisition of Pavlovian conditioned approach behavior to a food-paired cue.

#### Conditioned Reinforcement

Chronic semaglutide treatment enhanced the conditioned reinforcing properties of the lever-cue (**Figure 2e-g**). Although both treatment groups responded more in the active nose port that resulted in lever-cue presentation (Effect of Port (F_1,60_ = 147.62, *p* < 0.001)), SEMA-treated rats showed enhanced responding relative to VEH-treated rats (Treatment (F_1,60_ = 11.68, *p* = 0.001); Port x Treatment (F_1,60_ = 13.25, *p* < 0.001)) (**Figure 2e**). While SEMA-treated rats responded significantly more than VEH-treated rats in the active nose port (t = 3.834, *p* < 0.001; Cohen’s d = 0.959), there was no difference between groups in responding in the inactive nose port (t = 0.455, *p* = 0.632; Cohen’s d = 0.114). There was not a significant effect of Sex (F_1,60_ = 0.03, *p* = 0.864) on nose port responding, but there was a significant Port x Sex interaction (F_1,60_ = 9.80, *p* = 0.003). Males responded more in the inactive port than females (*p* = 0.003), with no sex differences in responding in the active port (*p* = 0.099). Upon lever presentation, SEMA-treated rats made more contacts with the lever than VEH-treated rats (Effect of Treatment (F_1,60_ = 4.22, *p* = 0.044)); post hoc t = 2.085, *p* = 0.041; Cohen’s d = 0.521) (**Figure 2f**). There was not a significant effect of Sex (F_1,60_ = 0.003, *p* = 0.956) nor a Treatment x Sex interaction (F_1,60_ = 0.16, *p* = 0.688) for lever-cue contacts during the conditioned reinforcement test. In agreement with the above, the incentive value index was greater for rats treated with semaglutide relative to those treated with vehicle (Effect of Treatment (F_1,60_ = 9.91, *p* = 0.003)); post hoc t = 3.145, *p* = 0.003; Cohen’s d = 0.786) (**Figure 2g**). There was not a significant effect of Sex (F_1,60_ = 2.04, *p* = 0.159) nor a Treatment x Sex interaction (F_1,60_ = 0.062, *p* = 0.803) for the incentive value index. Together, these data indicate that chronic semaglutide treatment enhances the incentive motivational value of a cue previously associated with food reward.

#### Progressive Ratio Responding

Chronic semaglutide treatment enhanced progressive ratio responding for the food reward (**Figure 2h-j**). Both groups of rats distinguished between the active and inactive ports, with more responding in the active port (Port (F_1,28_ = 51.99, *p* < 0.001)). However, relative to those treated with vehicle, rats treated with semaglutide showed more responding into the active port that resulted in reward delivery (Treatment (F_1,28_ = 7.43, *p* = 0.011); post hoc SEMA vs VEH, t = 2.835, *p* = 0.009; Cohen’s d = 1.002) (**Figure 2h**). There were no significant differences in responding in the inactive port (Port x Treatment interaction (F_1,28_ = 8.36, *p* = 0.007); posthoc SEMA vs VEH (t = 1.962, *p* = 0.067; Cohen’s d = 0.694)), and no effect of Sex (F_1,28_ = 1.25, *p* = 0.272). SEMA-treated rats also had significantly higher breakpoints (Treatment (F_1,28_ = 10.45, *p* = 0.003)); post hoc t = 3.256, *p* = 0.003; Cohen’s d = 1.151) (**Figure 2i**) and received more rewards in total (Treatment (F_1,28_ = 9.9, *p* = 0.004); post hoc t = 3.125, *p* = 0.004; Cohen’s d = 1.105) (**Figure 2j**) relative to VEH-treated rats. Contrary to our hypothesis, these findings indicate that rats treated with chronic semaglutide are more motivated to respond for small food rewards.

When analyzed across the entire population, responding in the active port during the progressive ratio test was significantly correlated with responding in the active port during conditioned reinforcement (r^2^ = 0.486, *p* = 0.005, *data not shown*). However, when analyzed within groups, responding in the active port during the progressive ratio and conditioned reinforcement tests was significantly correlated in VEH-treated rats (r^2^ = 0.701, *p* = 0.002), but not in SEMA-treated rats (r^2^ = 0.281, *p* = 0.292). Thus, we do not believe that greater responding at the active port during the conditioned reinforcement test impacted responding during the progressive ratio test for semaglutide-treated rats.

### Experiment 2

#### Weight Change

As in Experiment 1, chronic semaglutide treatment reduced weight gain in male rats and induced weight loss in female rats relative to vehicle-treated counterparts (**Figure 3a-b**). This is supported by significant main effects of Treatment (F_1,59_ = 149.17, *p* < 0.001), Day (F_16,944_ = 37.46, *p* < 0.001), and Sex (F_1,59_ = 88.61, *p* < 0.001) (**Figure 3a**). Additionally, there were significant Treatment x Day (F_16,944_ = 29.81, *p* < 0.001) and Day x Sex interactions (F_16,944_ = 29.02, *p* < 0.001). After Day 1 of treatment, SEMA-treated rats weighed significantly less than VEH-treated rats (regardless of sex) for the remainder of the experiment (*p* ≤ 0.004) (**Figure 3a**). At the end of the 17-day treatment regimen, both male and female rats treated with semaglutide had reduced weight gain with respect to baseline body weight relative to those treated with vehicle (**Figure 3b**) (Males: t = 9.218, *p* < 0.001; Cohen’s d = 3.313; Females: t = 6.808, *p* < 0.001; Cohen’s d = 2.412). There were significant main effects of Treatment (F_1,59_ = 131.38, *p* < 0.001) and Sex (F_1,59_ = 104.17, *p* < 0.001), and a Treatment x Sex interaction (F_1,59_ = 7.02, *p* = 0.010) on the final day of treatment. Similar to Experiment 1, most male rats treated with semaglutide showed attenuated weight gain while most female rats treated with semaglutide showed weight loss relative to baseline levels; but both sexes showed an approximate 10% difference in the change from baseline body weight relative to those treated with vehicle.

#### Pavlovian Conditioned Approach (PavCA)

Experiment 2 was designed to classify rats as sign-trackers, intermediate responders, or goal-trackers prior to administration of semaglutide or vehicle to assess potential phenotype-dependent effects. However, the distribution of phenotypes was skewed, with a predominance of sign-trackers and relatively few goal-trackers, limiting our ability to perform sufficiently powered statistical comparisons between groups. Therefore, for subsequent data analyses and illustrations, phenotype groups were collapsed.

Chronic semaglutide treatment had no effect on the expression of PavCA behavior. Table 2 presents the LMM analyses of the individual behavioral measures (i.e., lever contacts, probability of lever contact, latency to lever contact, magazine entries, probability of magazine entry, and latency to magazine entry) used to calculate PavCA Scores. These metrics are presented separately for Sessions 1-5 (pre-treatment, Acquisition) and 6-10 (during treatment, Expression). For behavior across Sessions 1-5, there was a significant main effect of Session (*p* ≤ 0.012) for each measure except for food-cup entries (F_4,72_ = 2.29, *p* = 0.068), indicative of a change in behavior as the cue-reward relationship was learned. There were no significant main effects of Treatment for any of the measures (*p* ≥ 0.723) across Sessions 1-5, as was expected given that Treatment groups were counterbalanced across these measures. Female rats overall showed a higher probability to contact the lever-cue (Effect of Sex: F_1,64_ = 6.08, *p* = 0.016). However, there were no main effects of Sex for any of the other measures (*p* ≥ 0.076). For Sessions 6-10, there was a significant main effect of Session for each measure (*p* ≤ 0.007), but no significant main effects of Treatment (*p* ≥ 0.184) or Sex (*p* ≥ 0.495).

As shown in **Figure 3c-d**, by the end of PavCA training for Sessions 1-5, all groups tended to sign-track (i.e., PavCA Score ≥ 0.5) rather than goal-track. Congruent with the findings described above for individual metrics, there was a significant main effect of Session (F_4,73_ = 7.94, *p* < 0.001), but no main effect of Treatment (F_1,73_ = 0.03, *p* = 0.870) or Sex (F_1,73_ = 3.16, *p* = 0.08), and no significant interactions (*p* ≥ 0.095) for PavCA Scores across Sessions 1-5 (**Figure 3c**). For Sessions 6-10, the tendency to sign-track increased with further training (Effect of Session: F_4,84_ = 9.28, *p* < 0.001), but the expression of PavCA behavior was not affected by chronic semaglutide treatment (Treatment (F_1,60_ = 0.00, *p* = 0.998)) or Sex (F_1,60_ = 0.04, *p* = 0.841). There were no significant differences between males and females across individual sessions (*p* ≥ 0.391), and no other interactions were statistically significant (*p* ≥ 0.084).

In agreement with the data above, there were no significant differences between treatment groups in the PavCA Index which was averaged across Sessions 4-5 (Acquisition) and 9-10 (Expression) (**Figure 3d**). There was no effect of Treatment (F_1,59_ = 0.03, *p* = 0.861), Sex (F_1,59_ = 0.82, *p* = 0.370), or Treatment x Sex interaction (F_1,59_ = 2.29, *p* = 0.136) for the Acquisition phase, nor were there significant effects for the Expression phase (Treatment (F_1,59_ = 0.09, *p* = 0.760), Sex (F_1,59_ = 0.01, *p* = 0.909), Treatment x Sex interaction (F_1,59_ = 3.25, *p* = 0.077)). Thus, consistent with the findings from Experiment 1, chronic semaglutide treatment does not appear to affect the expression of Pavlovian conditioned approach behavior to a food-paired cue.

#### Conditioned Reinforcement

As in Experiment 1, chronic semaglutide enhanced the conditioned reinforcing properties of the lever-cue (**Figure 3e-g**). Relative to vehicle, rats treated with semaglutide showed enhanced responding in the active nose port that resulted in lever-cue presentation, but not in the inactive port (**Figure 3e**; Effects of Port (F_1,59_ = 86.25, *p* < 0.001); Port x Treatment interaction (F_1,59_ = 4.12, *p* = 0.047); post hoc SEMA vs VEH active port: t = 2.124, *p* = 0.040; Cohen’s d = 0.535; post hoc SEMA vs VEH inactive port: t = 0.12, *p* = 0.885; Cohen’s d = 0.03). Overall, female rats made more nosepokes than male rats (Sex (F_1,59_ = 6.89, *p* = 0.011)), with responding in the active port primarily driving this effect (Port x Sex interaction (F_1,59_ = 10.22, *p* = 0.002); Sex effect at the active port (*p* = 0.003)). Congruent with greater responding in the active port, rats treated with semaglutide made more lever contacts than those treated with vehicle (**Figure 3f**; Effect of Treatment (F_1,59_ = 5.27, *p* = 0.025); post hoc t = 2.343, *p* = 0.022; Cohen’s d = 0.591), and SEMA-treated rats had a significantly higher Incentive Value Index than VEH-treated rats (**Figure 3g**; Treatment (F_1,59_ = 6.65, *p* = 0.012); post hoc t = 2.607, *p* = 0.011; Cohen’s d = 0.657). Although there were no sex differences for lever contacts, female rats had a significantly higher Incentive Value Index than males (Sex (F_1,59_ = 6.43, *p* = 0.014)), but there was no Treatment x Sex interaction (F_1,59_ = 0.019, *p* = 0.890). Together, these data align with those reported in Experiment 1, suggesting that chronic semaglutide treatment enhances the incentive motivational value of a cue previously associated with food reward.

#### Progressive Ratio Responding

Chronic semaglutide treatment enhanced progressive ratio responding for the food reward (**Figure 3h-j**), similar to Experiment 1. While both groups responded more in the active port than the inactive port (Effect of Port (F_1,59_ = 83.19, *p* < 0.001); posthoc (VEH: *p* = 0.011; SEMA: *p* < 0.001), SEMA-treated rats responded more in both ports compared to VEH-treated rats (Treatment (F_1,59_ = 28.11, *p* < 0.001); Port x Treatment interaction (F_1,59_ = 28.99, *p* < 0.001); posthoc active port: t = 4.984, *p* < 0.001; Cohen’s d = 1.256; post hoc inactive port: t = 3.545, *p* < 0.001; Cohen’s d = 0.893). Additionally, females made more nosepokes overall than males (Effect of Sex (F_1,59_ = 13.94, *p* < 0.001); Port x Sex interaction (F_1,59_ = 13.35, *p* < 0.001); posthoc Active: *p* < 0.001; Inactive: *p* = 0.004). In agreement with the nose port data, SEMA-treated rats had higher breakpoints relative to VEH-treated rats (**Figure 3i**; Effect of Treatment (F_1,59_ = 34.41, *p* < 0.001); post hoc t = 5.361, *p* < 0.001; Cohen’s d = 1.351), and female rats had higher breakpoints relative to male rats (Effect of Sex (F_1,59_ = 18.53, *p* < 0.001)); but treatment did not impact the sex effects (Treatment x Sex interaction (F_1,59_ = 1.16, *p* = 0.287)). Rats treated with chronic semaglutide also received more rewards than those treated with vehicle (**Figure 3j**; Effect of Treatment (F_1,59_ = 32.79, *p* < 0.001); t = 5.168, *p* < 0.001; Cohen’s d = 1.302), and females received more rewards than males (Effect of Sex (F_1,59_ = 21.97, *p* < 0.001)), but treatment did not impact the sex effects (Treatment x Sex interaction (F_1,59_ = 0.01, *p* = 0.922)). Consistent with Experiment 1, these findings indicate that chronic semaglutide treatment enhances the motivation to respond for small food rewards.

When analyzed across the entire population, responding in the active port during the progressive ratio test was significantly correlated with responding in the active port during conditioned reinforcement (r^2^ = 0.385, *p* = 0.002). However, when analyzed within groups, responding in the active port during the progressive ratio and conditioned reinforcement tests was not significantly correlated in VEH-treated rats (r^2^ = 0.301, *p* = 0.1) or SEMA-treated rats (r^2^ = 0.349, *p* = 0.05). Thus, we do not believe that responding during the conditioned reinforcement test impacted responding during the progressive ratio test.

### Experiment 3

#### Weight Change

As with Experiments 1 and 2, chronic semaglutide treatment reduced weight gain in males and induced weight loss in females relative to vehicle-treated controls (**Figure 4a-b**). This is supported by significant main effects of Treatment (F_1,28_ = 86.1, *p* < 0.001), Day (F_14,392_ = 84.4, *p* < 0.001), and Sex (F_1,28_ = 119.61, *p* < 0.001). Although there was not a significant Treatment x Sex interaction (F_1,28_ = 0.18, *p* = 0.678), the majority of female rats treated with semaglutide showed a decrease in body weight relative to baseline levels, whereas all male rats showed attenuated weight gain rather than weight loss. In support, at the end of the 17-day treatment regimen, both male and female rats treated with semaglutide had reduced weight gain with respect to baseline body weight relative to those treated with vehicle (**Figure 4b**) (Males: t = 5.815, *p* < 0.001; Cohen’s d = 2.907; Females: t = 3.491, *p* = 0.004; Cohen’s d = 1.746). There were significant main effects of Treatment (F_1,28_ = 43.33, *p* < 0.001) and Sex (F_1,28_ = 84.57, *p* < 0.001), but not a significant interaction on the final day of treatment. These results are consistent with Experiments 1 and 2, with chronic semaglutide causing a ≈10% or greater difference in the change from baseline body weight relative to those treated with vehicle.

**Figure 4.**
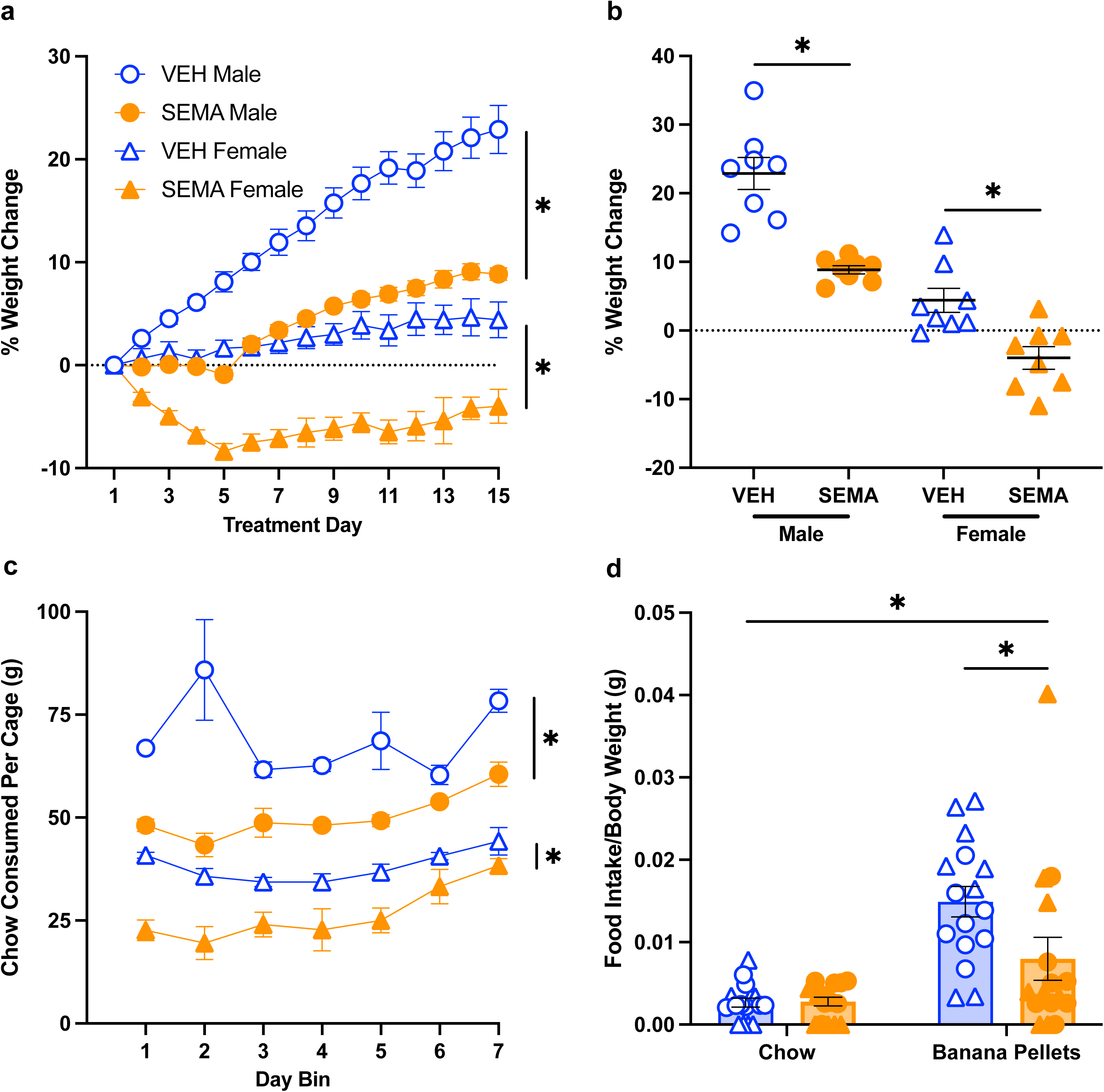
The effects of chronic semaglutide treatment on free consumption of chow and banana-flavored reward pellets. Data are shown for vehicle- (VEH, n = 16; 8 male and 8 female) and semaglutide- (SEMA, (n = 16; 8 male and 8 female) treated rats. (a) Mean ± SEM percent change in weight from baseline for VEH- and SEMA-treated rats across 15 days of treatment. Relative to vehicle, chronic semaglutide treatment reduced weight gain in males and induced weight loss in females. (b) Black bars show mean ± SEM weight change from baseline following the final treatment on Day 15, and each data point represents an individual rat. Semaglutide treatment reduced weight gain in both males and females. (c) Mean ± SEM chow consumed per cage of VEH- and SEMA-treated rats presented in 2-day bins over the course of Experiment 3. Chronic semaglutide treatment reduced homecage chow consumption in males and females. (d) Mean ± SEM amount of chow or banana pellets consumed divided by body weight for VEH- and SEMA-treated rats during feeding tests. Each data point represents an individual rat. Relative to vehicle, chronic semaglutide treatment reduced the amount of banana pellets, but not chow, consumed during the free-feeding consumption test. \**p* < 0.05

#### Chow Consumption

Chronic semaglutide treatment reduced homecage chow consumption across a 14-day treatment period (**Figure 4c**; Effects of Treatment (F_1,12_ = 50.14, *p* < 0.001); Sex (F_1,12_ = 161.53, *p* < 0.001); Bin (F_6,72_ = 8.37, *p* < 0.001)). There was a significant Treatment x Bin interaction (F_6,72_ = 5.26, *p* < 0.001) and post hoc comparisons showed that SEMA-treated rats ate less chow than VEH-treated rats during each 2-day bin (*p* ≤ 0.012). No other interactions were statistically significant (*p* ≥ 0.087). These data further highlight the validity of our chronic semaglutide treatment regimen, showing that it attenuates homecage chow consumption.

#### Free-feeding Consumption Tests

During the free-feeding consumption tests, chronic semaglutide treatment reduced the amount of banana-flavored reward pellets consumed relative to vehicle but had no effect on chow consumption (**Figure 4d**). Overall, rats ate more banana-flavored pellets than chow (main effect of Food Type (F_1,28_ = 32.27, *p* < 0.001; Treatment x Food Type interaction (F_1,28_ = 5.29, *p* = 0.029)). Post hoc comparisons showed that SEMA-treated rats ate significantly fewer banana pellets than VEH-treated rats (t = 2.167, *p* = 0.038; Cohen’s d = 0.766), with no difference between groups with respect to chow consumption (t = 0.17, *p* = 0.865; Cohen’s d = 0.060). No other interactions were statistically significant (*p* ≥ 0.058) for the free-feeding consumption test. Thus, chronic semaglutide treatment appears to decrease the consumption of salient food rewards when such rewards are unlimited.

## Discussion

Contrary to our hypothesis, we did not observe effects of chronic semaglutide administration on the acquisition or expression of Pavlovian conditioned approach behavior. Furthermore, opposite to what we had anticipated, chronic semaglutide treatment *enhanced* the conditioned reinforcing properties of a food-associated cue as well as progressive ratio responding for a small food reward. Notably, these effects were consistent across both Experiments 1 and 2. We also found that chronic semaglutide treatment reduces homecage chow consumption and the consumption of banana-flavored reward pellets when access is unlimited. Together, these findings suggest that chronic semaglutide acts on incentive motivational processes to enhance the “potency”—or amplify the reinforcing effects—of small food rewards and associated cues, resulting in quicker satiety and reduced consumption when food is readily available.

The enhanced responding observed on the conditioned reinforcement test suggests that chronic semaglutide treatment increases the incentive value of the cue previously associated with food reward (Robinson & Flagel, 2009). However, we did not observe corresponding enhancements in sign-tracking behavior during Pavlovian conditioning, which is also indicative of increased incentive value attributed to the food cue (Robinson & Flagel, 2009). It is possible that effects of semaglutide on Pavlovian conditioned approach behavior were masked, as most rats in these experiments predominantly exhibited sign-tracking behavior. That is, we were unable to further enhance their attraction to the food cue, regardless of whether semaglutide was administered prior to learning the cue-reward relationship (Experiment 1) or after their conditioned response had already been acquired (Experiment 2). Additionally, during each Pavlovian conditioned approach session, the lever-cue was available for only 8 s per trial across 25 trials, resulting in approximately 200 s of availability per session. It is possible, therefore, that semaglutide altered the attribution of incentive value to the lever-cue, but these effects may not have been detected due to limited cue access during the Pavlovian conditioned approach paradigm. By comparison, during the conditioned reinforcement test, the lever is presented in 2-second intervals following a response in the active nose port, resulting in an average of approximately 300 seconds of total lever availability per session. Thus, any facilitative effects of semaglutide on incentive salience attribution during Pavlovian conditioning may also have been masked by the restricted access to the lever-cue.

The apparent discrepancy between the effects of chronic semaglutide on the conditioned reinforcing properties of the food-associated cue and its lack of impact on cue-elicited Pavlovian conditioned approach behavior may reflect the involvement of distinct neural systems underlying these behaviors. Indeed, separate neurobiological mechanisms are known to govern Pavlovian and instrumental learning (Balleine, 2005; Todd et al., 2014; Bouton et al., 2021). Notably, the conditioned reinforcement paradigm employed in our study required rats to learn a novel instrumental response for cue presentation and was conducted under extinction conditions, with no food reward provided during training. Further investigation is warranted to elucidate how semaglutide interacts with these distinct learning systems and to determine the conditions under which its behavioral effects are most pronounced.

The effects observed on progressive ratio responding following chronic semaglutide administration were unexpected yet paralleled the conditioned reinforcement results, supporting the idea that our chronic treatment regimen enhanced the motivational value of both the food reward and its associated cue. When rats responded for the small food reward (one 45-mg banana-flavored pellet), they continued to find it reinforcing; semaglutide-treated rats worked for and consumed an average of approximately 25 pellets (1.125 g) during the progressive ratio test, while vehicle-treated rats consumed an average of approximately 15 pellets (0.675 g). When given free access to the same reward pellets, chronic semaglutide-treated rats consumed an average of 44 pellets (2 g), whereas vehicle-treated rats consumed more than 2.5 times that amount—an average of 111 pellets (5 g). This pronounced difference suggests that chronic semaglutide treatment enhances motivated responding for food reward up to a certain limit and this limit was not reached during the progressive ratio test but was during the free-access test. The decrease in homecage chow consumption following chronic semaglutide treatment further supports the idea that semaglutide may selectively enhance motivational responding for small, palatable food rewards while simultaneously reducing overall food intake. These findings are consistent with prior preclinical studies demonstrating that chronic semaglutide reduces meal size without affecting meal frequency in the homecage (Cawthon et al., 2023). Collectively, these results bolster our conclusion that semaglutide enhances the “potency” of consumed rewards and thereby promotes earlier satiety.

Our findings differ from much of the existing preclinical literature examining the effects of GLP-1 receptor agonists on food-motivated responding. For example, Exendin-4 has been shown to reduce progressive ratio responding and conditioned place preference for food reward (Dickson et al., 2012; Richard et al., 2016; Colvin et al., 2020). It should be noted, however, that prior studies primarily evaluated the effects of single or a few repeated injections, administered systemically or directly into the brain (Dickson et al., 2012; Richard et al., 2016; Vallӧf et al., 2016; Colvin et al., 2020). By contrast, our study implemented a chronic systemic treatment regimen, similar to that used by Cawthon et al. (2023), to more accurately model the clinical application of semaglutide in humans (Chao et al., 2023).

Cawthon et al. (2023) observed an enhancement, rather than a reduction, in 4% liquid sucrose consumption in rats treated with chronic semaglutide compared to those given vehicle. While the direction of this effect aligns with our findings, direct comparison is complicated, as we observed semaglutide-induced enhancement of food reward in the progressive ratio test, yet reduction in food intake during the free-access test. In the Cawthon et al. (2023) study, sucrose preference was assessed using a two-bottle test in the rats’ homecage, where food was also freely available. Furthermore, rats in their study were notably older and had higher baseline body weights than those used in our experiments. Nevertheless, we believe that using a chronic treatment regimen analogous to clinical practice provides important insights into the psychological and neural effects of GLP-1 receptor agonists and enhances the translational value of our findings.

The brain mechanisms underlying the effects of chronic semaglutide administration remain poorly understood. GLP-1 receptors are expressed throughout multiple brain regions, including the ventral tegmental area, nucleus accumbens, several hypothalamic nuclei, and the paraventricular nucleus of the thalamus (Gabery et al., 2020; Gӧke et al., 1995; Heppner et al., 2015; Jensen et al., 2018) — all of which play key roles in the attribution of incentive motivational value to rewards and associated cues (Haight & Flagel, 2014; Iglesias & Flagel, 2021; Flagel et al., 2011; Flagel & Robinson, 2017). Although semaglutide is capable of accessing the brainstem, septal nucleus, and hypothalamus, evidence suggests that it does not effectively cross the blood-brain barrier (Gabery et al., 2020). Instead, its central actions may occur via circumventricular organs and regions adjacent to the ventricles.

Reductions in reward-seeking behaviors mediated by GLP-1 receptor signaling are thought to involve dopaminergic pathways that are critical for incentive motivation (Iglesias et al., 2023; Flagel et al., 2011; Saunders et al., 2013; Berridge, 2007; Berridge & Robinson, 1998). GLP-1 analogs targeting midbrain dopamine neurons in the ventral tegmental area have been shown to attenuate high-fat food intake and decrease the excitatory synaptic strength of projections to the nucleus accumbens shell (Wang et al., 2015). Additionally, central administration of GLP-1 receptor agonists reduces sucrose-seeking behavior and dampens cue-evoked dopamine activity (Konanur et al., 2020).

Other neural sites implicated in appetitive behavior, such as the lateral hypothalamus (López-Ferreras et al., 2018a), supramammillary nucleus (López-Ferreras et al., 2019), and paraventricular nucleus of the thalamus (Ong et al., 2017), also mediate the effects of GLP-1 receptor agonists on both food intake and motivational drive. For example, activation of GLP-1 receptors in the paraventricular nucleus of the thalamus decreases neuronal excitability of projections to the nucleus accumbens (Ong et al., 2017). This same pathway encodes the incentive value of food-associated cues, with engagement to a greater degree in sign-trackers relative to goal-trackers (Haight et al., 2017). Ongoing work in our laboratory aims to further clarify the role of dopamine and the paraventricular nucleus of the thalamus-to-nucleus accumbens circuit in mediating the effects of chronic semaglutide observed in this study.

Importantly, our studies were conducted in non-obese animals, which may differ substantially from clinically obese populations with respect to metabolic and neural responses. Nevertheless, the present findings may have translational relevance for humans, given that they mirror clinical observations that semaglutide reduces food intake and supports weight management, potentially by modulating neural circuits involved in motivation and reward. Additional research is needed to determine whether the effects observed here generalize to models of obesity and to broader human populations.

In summary, our results demonstrate that chronic semaglutide administration amplifies the incentive salience of cues previously associated with food rewards and enhances motivational drive for food when access is limited. In contrast, under conditions of unrestricted food availability, semaglutide mitigates excessive consumption. This dual action suggests that semaglutide may intensify motivation for rewards in resource-constrained environments, while concurrently promoting restraint when abundance could otherwise trigger overconsumption. Elucidating the neural mechanisms underlying these effects presents a promising avenue for future research.

## Acknowledgments

The authors would like to thank members of the Flagel Laboratory for their support and insight regarding this research.

