## Supplementary Information for "Chronic semaglutide treatment enhances the incentive motivational value of a small food reward and associated cue in male and female rats"

**SUPPLMENTARY INFORMATION**

### Locomotor Response to Novelty

Following the last day of semaglutide (or vehicle) administration (i.e., the day following the progressive ratio test), rats were tested for locomotor response to novelty, an index of “sensation-seeking” that has been associated with vulnerability to addiction-like behaviors (Piazza et al., 1989; Flagel et al., 2014). Rats were taken to an unfamiliar testing room and placed in a novel cage made of clear acrylic (43 x 21.5 x 25.5 cm (high) and equipped with infrared photocell emitters mounted 2.3 and 6.5 cm above a metal grid floor. Cages were on racks separated by black acrylic walls. Rats were allowed to move freely around the cage, and infrared beam detectors recorded lateral and rearing movements upon beam deflection. The total number of lateral and rearing movements were then combined to create a cumulative activity score for each rat. This session lasted for 30 minutes.

*Data Analysis*

For both Experiments 1 and 2, cumulative locomotor activity was analyzed using ANOVAs with factors of Treatment (VEH vs. SEMA) and Sex (Male vs. Female). The significance level was $p < 0.05$ for all tests.

### Results

*Experiment 1*

The 32 rats (16 male, 16 female) from round 2 of Experiment 1 that underwent testing for progressive ratio (PR) responding, were also tested for locomotor response to novelty.

*Experiment 2*

All of the rats included in Experiment 2 (n=64; 32 male, 32 female) were tested for locomotor response to novelty.

These findings suggest that chronic semaglutide administration does not affect “sensation-seeking” behavior.
